# Evaluating Bacterial Viability in Faecal Microbiota Transplantation: A Comparative Analysis of In Vitro Cultivation and Membrane Integrity Methods

**DOI:** 10.1101/2024.02.08.579472

**Authors:** Ivana Cibulková, Veronika Řehořová, Marek Wilhelm, Hana Soukupová, Jan Hajer, František Duška, Helena Daňková, Monika Cahová

**Author notes:** Address correspondence to (M.C.).

## Abstract

**Background:** Faecal microbiota transplantation (FMT) is a developing therapy for disorders related to gut dysbiosis. Despite its growing application, standardized protocols for FMT filtrate preparation and quality assessment remain undeveloped. The viability of bacteria in the filtrate is crucial for FMT’s efficacy and for validating protocol execution. We compared two methods—in vitro cultivation and membrane integrity assessment—for their accuracy, reproducibility, and clinical applicability in measuring bacterial viability in frozen FMT stool filtrate.

**Methods:** Bacterial viability in stool filtrate was evaluated using (i) membrane integrity through fluorescent DNA staining with SYTO9 and propidium iodide, followed by flow cytometry; and (ii) culturable bacteria counts (colony-forming units, CFU) under aerobic or anaerobic conditions.

**Results:** We refined the bacterial DNA staining protocol integrated with flow cytometry for stool samples. Both the membrane integrity-based and cultivation-based methods exhibited significant variability in bacterial viability across different FMT filtrates, without correlation. The cultivation-based method showed a mean coefficient of variance of 17%, ranging from 5.3% to 52.9%. Conversely, the membrane integrity approach yielded highly reproducible results, with a median coefficient of variance for viable cells of 0.9%, ranging from 8.5% to 0.04%.

**Conclusion:** Bacterial viability assessment using cultivation-dependent methods produces inconsistent outcomes. In contrast, the membrane integrity method offers robust and precise data, making it a viable option for routine faecal material evaluation in FMT.

## Introduction

Recent advancements in microbiome research have dramatically altered our perception of microorganisms’ role in human health. Historically, the human body’s resident bacteria were deemed foes, prompting strategies to shield ourselves from these and other microbiome entities. The technological progress driven by the Human Genome Project opened new possibilities in many research areas and, among others; it enabled the revealing of the enormous richness of the microbial world associated with human organism. This led to the shift in the perception of a multicellular eukaryotic organism defined solely or predominantly by its own genome to the idea of a holobiont, composed of the eukaryotic “macro”organism accompanied by a highly variable resident microbiota (the bacteria, archaea, viruses, and fungi) (1). Along with this change in paradigm there are various efforts to modulate the microbiome through dietary changes, prebiotics, probiotics, and faecal microbiota transplantation (FMT), the latter showing notable promise in altering gut microbiota composition significantly (1–3).

At present, FMT is officially recognized for treating recurrent *Clostridioides difficile* infection (CDI) in the US and EU (4–7). Beyond CDI, FMT’s efficacy extends to conditions like hepatic encephalopathy, inflammatory bowel disease (particularly ulcerative colitis), irritable bowel syndrome, neurological disorders, and the transmission of multidrug-resistant organisms (8–10). Over 460 clinical trials are underway, exploring its potential across various health issues, including obesity and multiple sclerosis (11). In 2019, Europe reported 1874 hospital-based FMT procedures for both routine and research purposes (10). Despite its established role, particularly in CDI, standardized guidelines for FMT application are lacking (12). International consensus offers best practice recommendations (8,13,14); however, protocols for donor screening, filtrate preparation, and management differ widely, often without transparency between institutions (10). Notably, guidelines on FMT filtrate preparation and stool filtrate quality, especially regarding bacterial viability, remain undefined. Evidence suggests that higher bacterial viability correlates with improved FMT outcomes (8,15,16).

Regulatory perspectives on FMT vary globally; some nations regulate the stool product as a pharmaceutical, while others classify it alongside human cell and tissue products. In the USA, Canada, France, and the Czech Republic, FMT filtrate is considered an investigational drug, necessitating standardization for safety and efficacy (17). Bacterial viability is a key characteristic influencing both therapeutic effectiveness and protocol adherence (15,18). Thus, identifying a straightforward, dependable, and cost-effective method for assessing bacterial viability is imperative for FMT centers and stool banks to function effectively and within regulatory framework.

This study aims to establish a precise and practical approach for assessing bacterial viability in stool filtrate, facilitating its integration into standard operating procedures by FMT centers and biobanks.

## Materials and Methods

### Donor Characteristics

Human biological samples were collected during the FEBATRICE study, approved by the Ethics Committee for Multi-Centric Clinical Trials of FNKV University Hospital (Decision No. KH/40/00/2021). Donors underwent screening as per the FEBATRICE clinical trial protocol (NCT05430269). Altogether 13 stool samples were obtained from nine donors (2 females, 7 males), median age 36 yrs (min 25; max 42), median BMI 23.7 (min 17.6; max 27.8). The study protocol is in accordance with the Declaration of Helsinki (2013 amendment) and all the donors gave a written, prospective informed consent.

### Human Stool Processing

Stool from donors, devoid of diarrhea and free from mucus or blood, was processed within four hours post-defecation following the standard operating procedure (19). Samples were diluted in sterile 0.9% NaCl, homogenized with a dedicated hand blender (Mulinex), filtered for uniformity, mixed with 10% glycerol by volume, and stored at -80°C for up to one year. The sample handling pipeline is shown in Figure 1.

**Figure 1:**
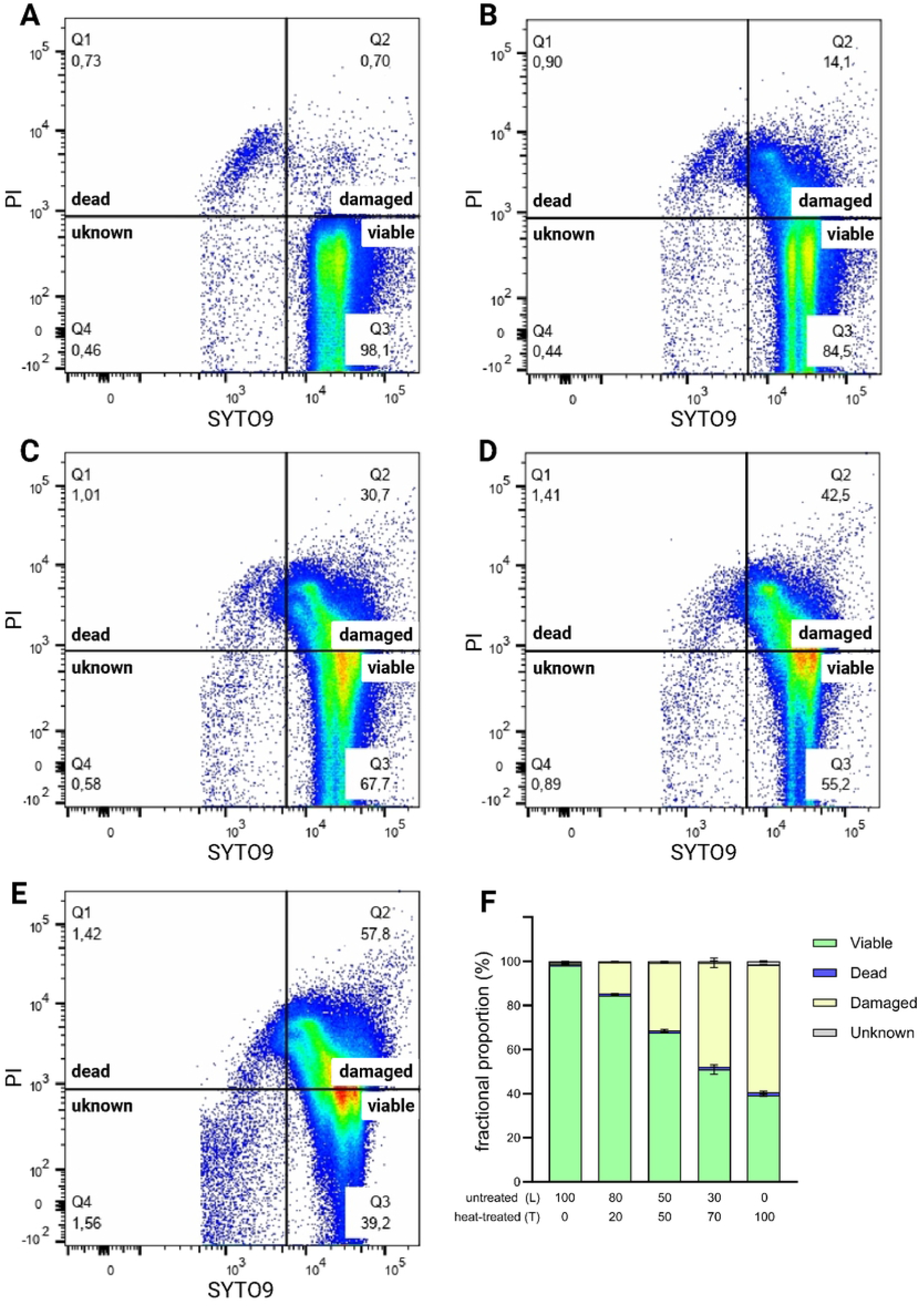
Working pipeline for the preparation, storage and viability assessment of FMT stool filtrate samples by flow cytometry and cultivation

### Mice Stool Sample Processing

The stool samples from germ-free (GF) C57Bl6/J mice were obtained from the colony bred at the Gnotobiology laboratory Institute of Microbiology of the CAS, Novy Hradek, CR and treated in sterile way. The samples from conventionally bred (CONV) C57Bl6/J mice were obtained from the breeding facility IKEM, Prague. All samples were kept in -20°C. Prior the experiment, mouse feces (100mg) were thawed at room temperature for 10 min, diluted in 0.9% NaCl 1:4 (wt/wt), homogenized by vigorous vortexing and glycerol (10% wt/wt) was added.

### Viability Determination in Pure Bacterial Culture

Overnight culture of E. coli ATCC 10536 (CCM 3988) (50 ml of LB medium, 37°C, 200 RPM) was harvested by centrifugation. The pellet was resuspended in 10 mL 0.9% NaCl solution and then diluted by the decimal series to a final concentration of 1:20000. One half of the sample was left (live bacteria) and the other half was treated at 70 °C for 10 minutes (heat-treated bacteria). Before staining, mixtures with different proportions of live and heat-treated cells were prepared.

### Viability Determination in Human Stool Samples

Frozen FMT filtrates were thawed at 36°C for 10 minutes and diluted by the decimal series to a final concentration of 1:10000. Wider-necked tips were used to prevent clogging due to solid particles in the sample. If required, the samples were pretreated by incubation at 80°C for 15 minutes. In some experiments, the dilution and staining procedure was performed under semianaerobic conditions. The saline solution was dispensed á 9 ml into sterile glass vials, bubbled with nitrogen for 15 min and sealed with a rubber stopper with an aluminum rim. The FMT filtrate prepared in the standard manner was thawed (closed vessel, 10 min, 36°C). After thawing, the vessel was opened and the content drawn with a 18G needle into a syringe and 1 ml of the suspension was injected with a needle through the rubber cap into a vial with degassed saline (dilution 1:10). The content of the vial was vortexed and the same process was repeated until we reached a dilution of 1:10 000. An aliquot (500 ul) of the diluted suspension was transferred to another tube and the stains (SYTO9 and PI) were added. The tube was placed in a 50 ml falcon which was inflated with N2 gas, closed with a screw cap and sealed with a parafilm. The sample was stored in this way until measured on a flow cytometer. The aerobic and anaerobic dilution scheme is shown in Figure 2.

**Figure 2:**
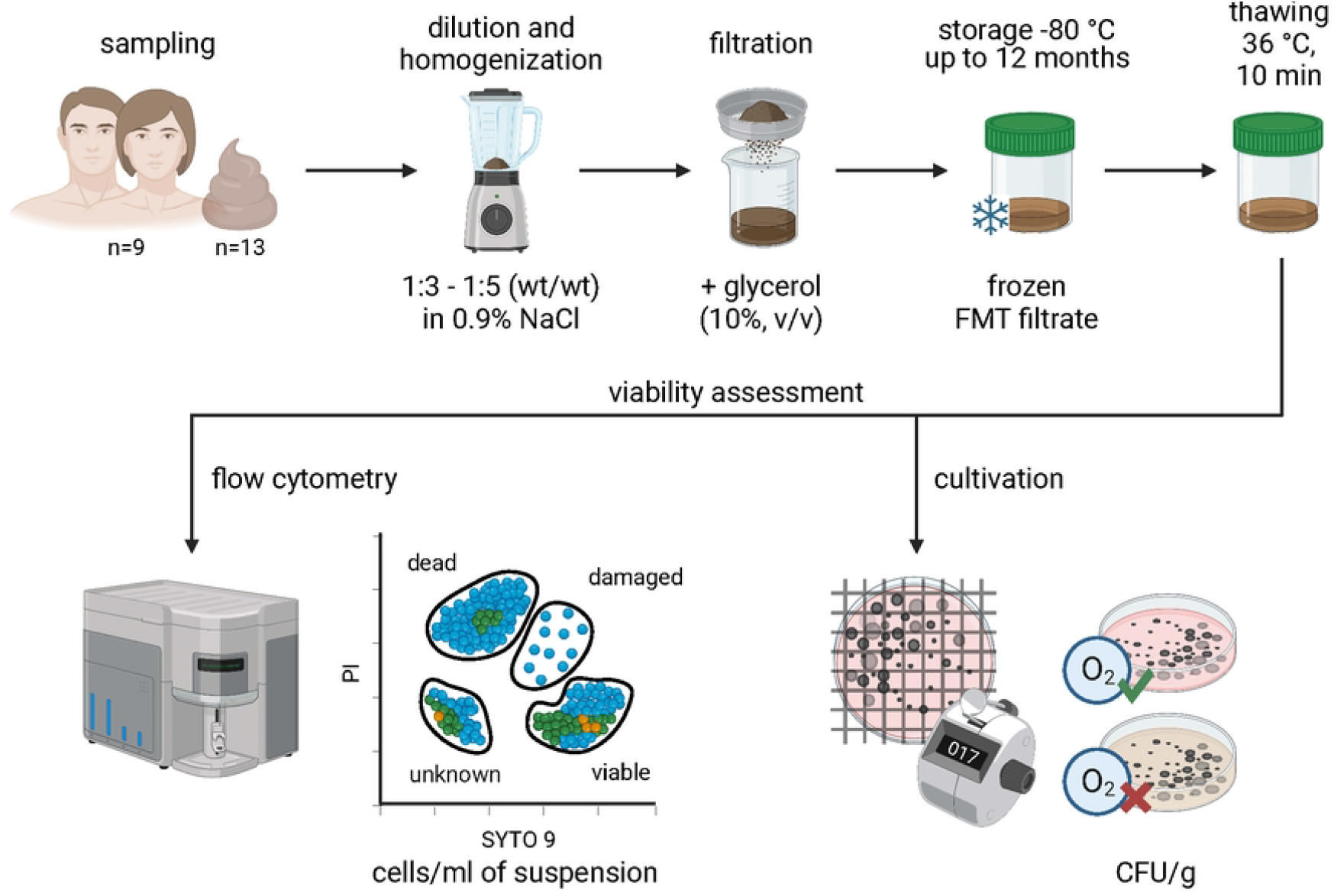
Procedure for aerobic and semi-anaerobic staining of FMT stool filtrate samples for the assessment of bacterial viability using flow cytometry.

### Flow Cytometry

LIVE/DEAD™ *Bac*Light™ Bacterial Viability Kit, for microscopy & quantitative assays, catalog number L7012 was performed to determine the viability of bacteria in faecal samples according to manufacturer instructions. Briefly, 500 μL of mixture, 0.75μL of SYTO9 and 0.75 μL of propidium iodide (PI) were mixed together and incubated for 15 min in the dark. Total volume 100 μL of diluted sample were added to a flow cytometry analysis tube. All samples were analyzed in triplicate on a 4-laser BD LSR II flow cytometer with the excitation 488nm and emission 525/50nm (=500-550nm) for SyTO9 and 561nm excitation and 610/20nm emission (=600-620nm) for propidium iodide.

### Cultivation of Stool Microbiota

Thirteen FMT filtrates were cultured aerobically and anaerobically. Colony counts were translated into CFUs per gram of faecal filtrate. In the preliminary experiments (data not shown), we established the optimal dilution and culture methodology. Our finalized protocol involved taking 300 μl (0.3 ml) of the homogenate, which was subsequently measured for weight to verify the accuracy of the pipette sampling, and diluting it in 3 ml of normal saline. This dilution process was repeated four more times until the final dilution of 1:10000. The diluted aliquots were inoculated onto blood agar for aerobic cultivation and Schaedler agar for anaerobic cultivation. The blood agar plates were incubated for 24 hours for standard aerobic cultivation, while the Schaedler agar plates were incubated for 48 hours for standard anaerobic cultivation. The number of colony-forming units (CFUs) was counted and adjusted relative to 1 g of the original homogenate. All tests were performed in triplicates.

### Statistical Analysis

Statistical analysis was performed using the Mann–Whitney t-test to identify statistically significant differences between different protocols (Prism v5.0, GraphPad). Descriptive statistics were calculated and reported as means and standard deviations. A P < 0.05 was considered statistically significant.

## Results

### Optimization of Staining Procedure for Stool Samples

Kits designed for assessing bacterial viability are typically validated for in vitro-grown bacterial cultures. Given that stool samples constitute a markedly different matrix, we tailored the testing protocol. Establishing a reliable gating strategy necessitated differentiating bacterial cells from sample debris and background noise. First, we established a baseline signal using blank sample with deionized water and both fluorescent dyes to depict and quantify false-positive events caused by noise, impurities and aggregates in assay reagents. Figure 3 shows the blank baseline of “reagents only” fingerprint where deionized water is in place of samples. We clearly show and quantify the fraction of false-positive events being processed in each sample due to the presence of noise in the optical and electronic system, impurities in deionized water and aggregates in SYTO9 and PI reagents which are present in all samples at equal quantities.

**Figure 3:**
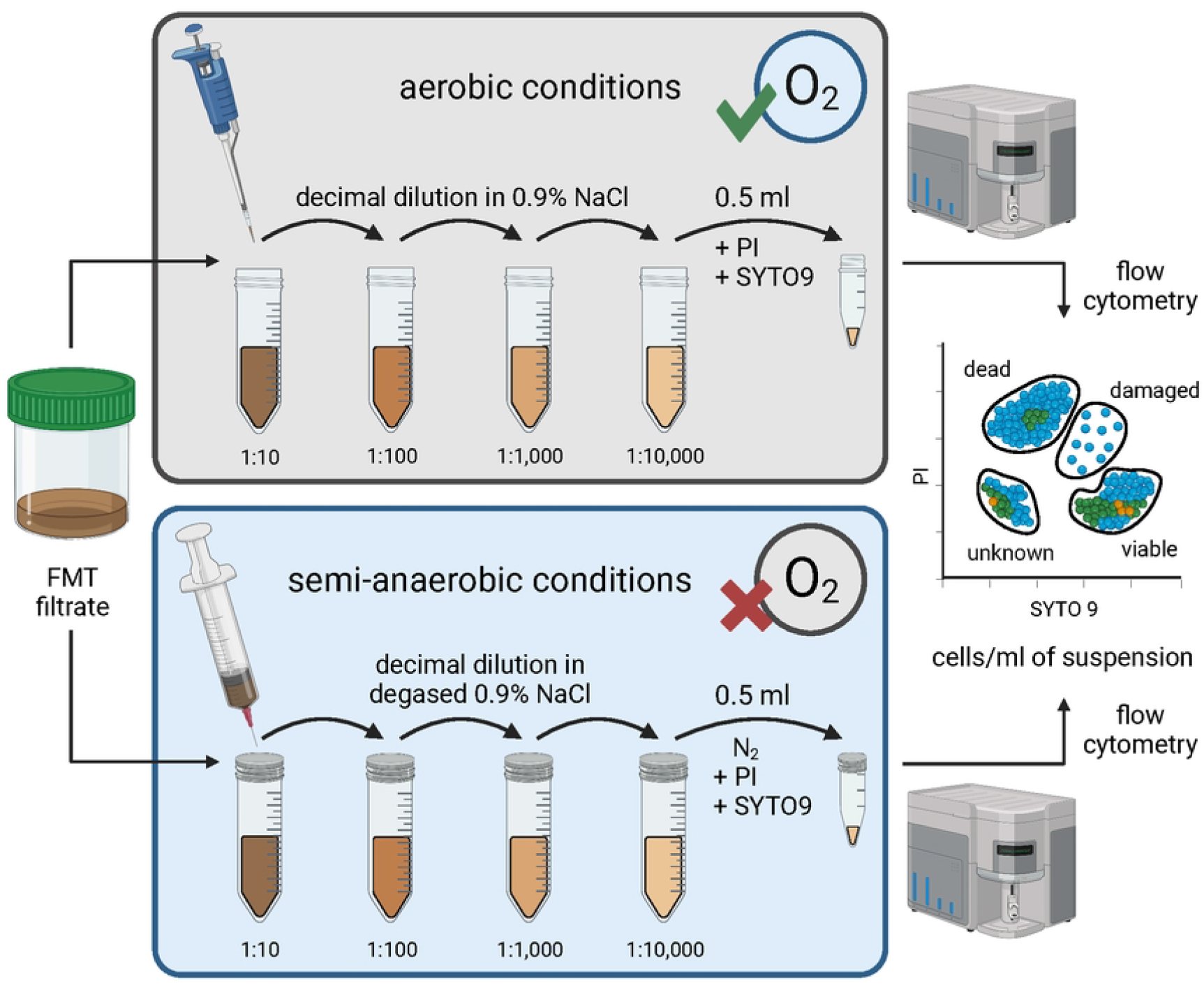
Baseline signal determination using blank sample with deionized water and both fluorescent dyes.

Stool samples inherently contain heterogeneous, autofluorescent components, such as bile derivatives, not accounted for in the standard staining methods. We used stool from germ-free (GF) mice, devoid of bacteria, to establish a noise threshold and compared this with stool from conventionally bred (CONV) mice, containing both bacterial and non-bacterial elements (Figure 4). This approach allowed us to define four viability quadrants based on SYTO9 and PI fluorescence, categorizing cells as dead, dying or damaged, viable, or of unknown status (20,21). We observed significantly fewer fluorescent events in GF mouse stool, attributing this to non-bacterial autofluorescence or staining artifacts.

**Figure 4:**
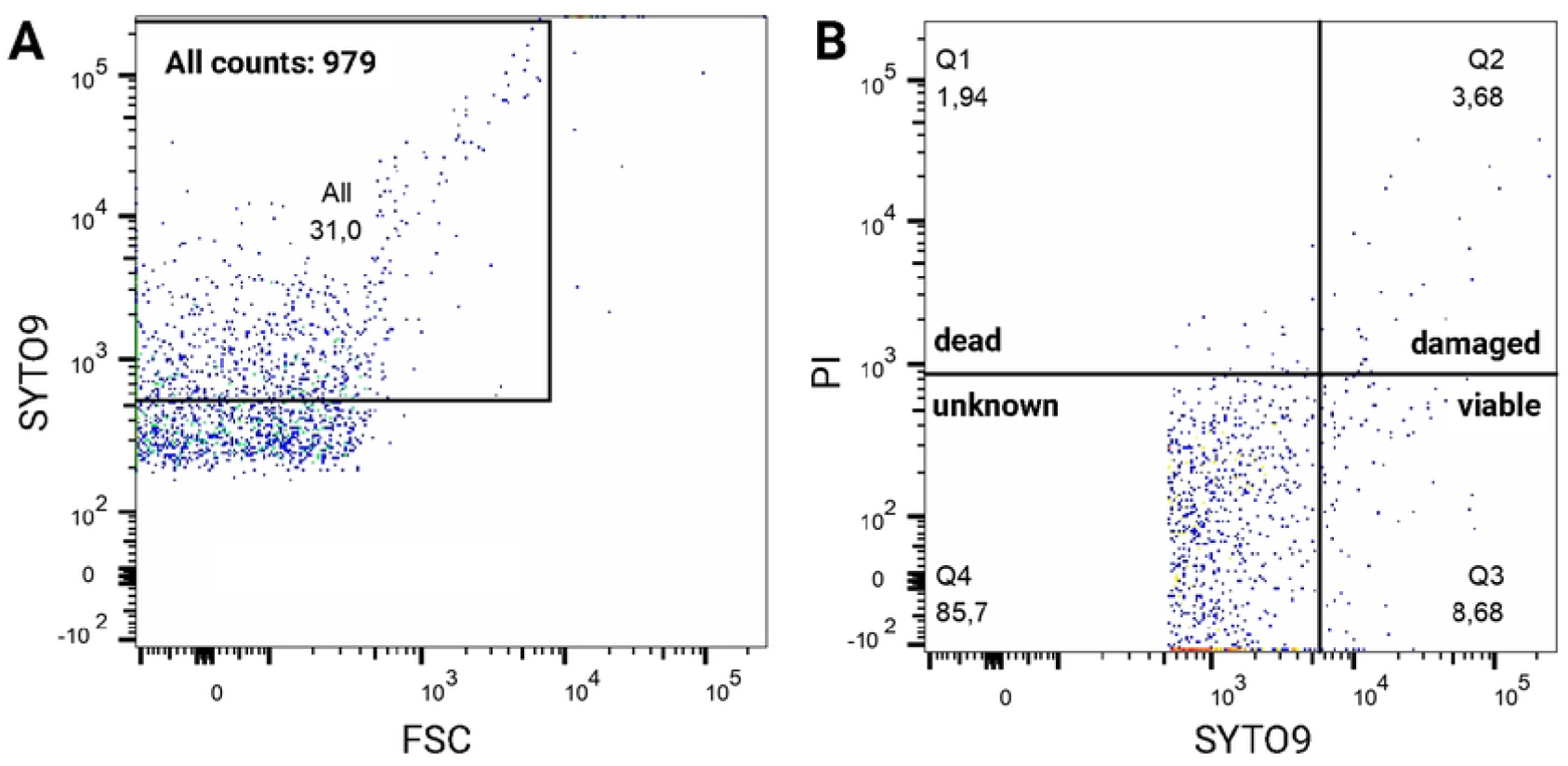
Optimization of the gating strategy for faecal samples. A,B: germ-free mice; C,D: conventional mice; E: quantification of the signal (absolute counts).

### Reliability of Viability Discrimination

To confirm this gating strategy and the accuracy of this assay, we analyzed mixtures of live (L) and heat-treated (T) *E. coli* cultures mixed in different ratios. The results demonstrated a proportional increase in non-viable cells with increased proportion of heat-treated bacteria, affirming the method’s ability to distinguish between viable and non-viable cells (Figure 5). The consistency of our method was evaluated also on stool samples, comparing native and heat-pre-treated samples (Figure 6). Similarly to pure bacterial culture, the percentage of live bacterial cells was significantly reduced by heat treatment (53.5± 0.7 vs. 1.4±0.1), while the percentage of dead bacteria increased accordingly (21.1± 0.8 vs. 68±1.4) with a low coefficient of variance (CV) across measurements, indicating that flow cytometry is a reliable tool for assessing faecal transplant viability.

**Figure 5:**
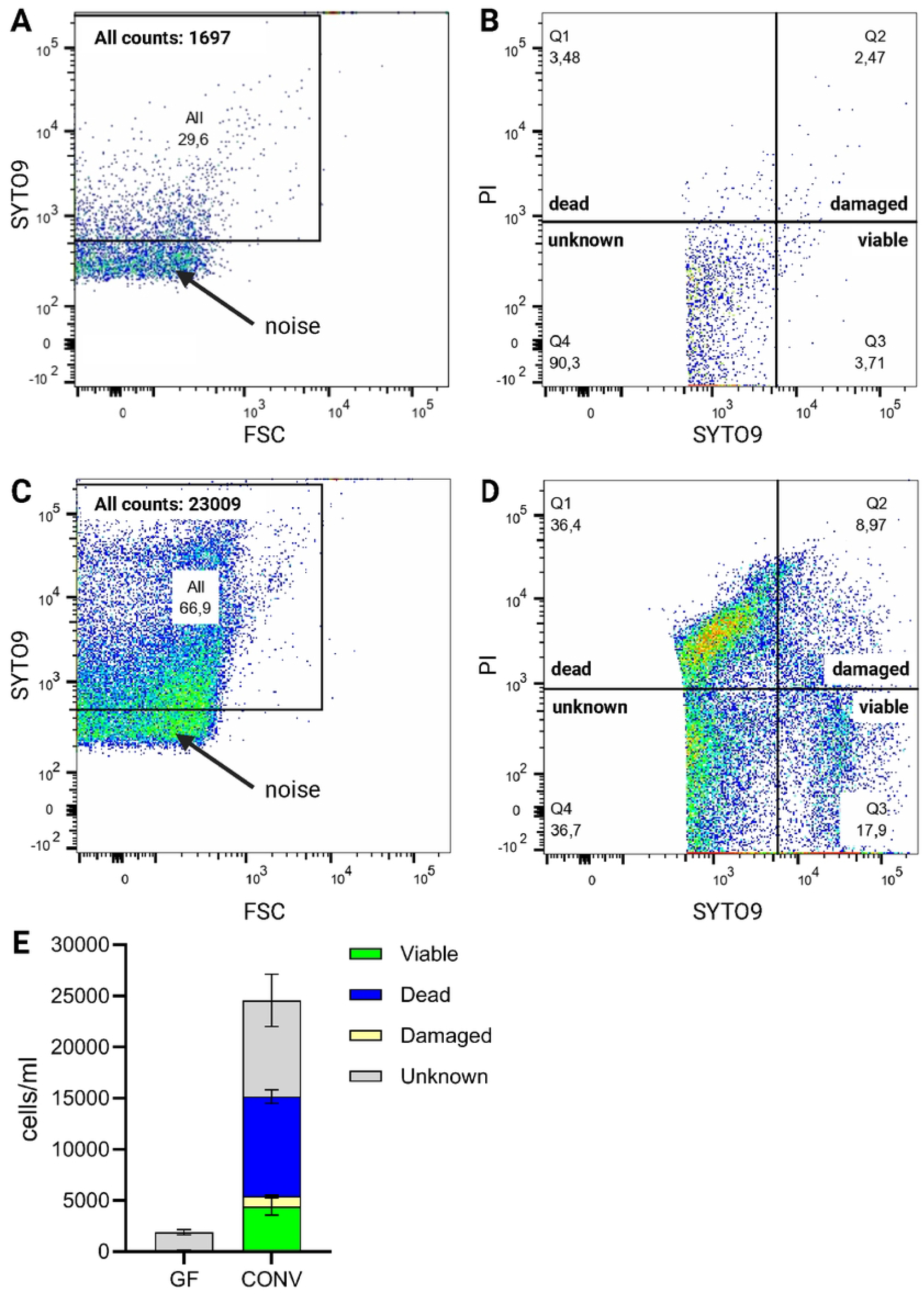
Quantification of live and dead cells in the mixture of untreated live (L) and heat-treated (T) culture of E. coli. Heat-treated samples were incubated for 10 minutes at 70 °C. A: untreated sample; B: L/T mixture ratio 80/20, C: L/T mixture ratio 50/50, D: L/T mixture ratio 30/70, E: L/T mixture ratio 0/100, F: quantification of the signal.

**Figure 6:**
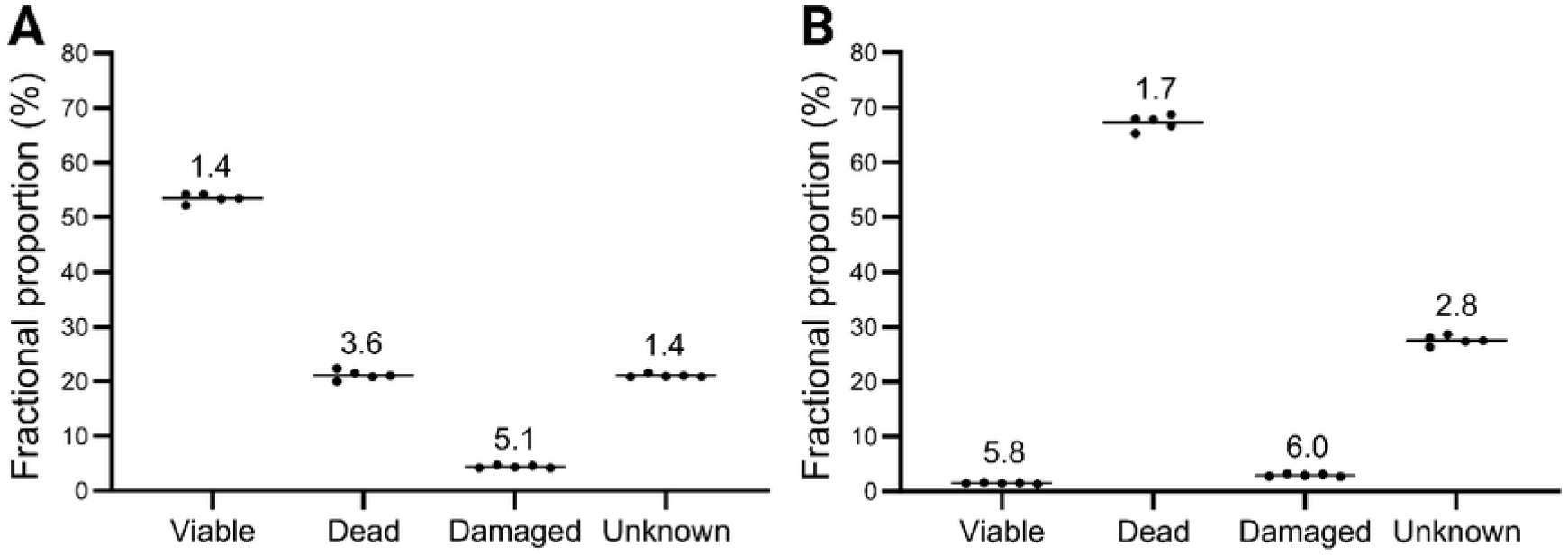
The viability of bacteria in human FMT stool filtrate sample. **A**: Thawed sample. **B**: Thawed and heat-treated sample. Heat-treated samples were incubated 10 min at 80°C. Each measurement was performed in five technical replicates. The lines represent the medians, the numbers indicate the median CV (%).

### Impact of Staining Conditions on Bacterial Viability

A significant proportion of intestinal bacteria are anaerobic and preparing the sample for analysis (approximately 2 hours) on the laboratory bench could reduce the viability of the anaerobic bacteria and significantly bias the results. To assess the effect of oxygen atmosphere during sample handling, we used two frozen aliquots from the same human donor and performed the staining procedure under both semi-anaerobic and aerobic conditions. The number of viable bacteria decreased by 8% under aerobic compared to semi-anaerobic conditions (49.6±5.3 versus 57.3±5.5%). The percentage of dead cells (25.2±7.8 vs. 13.1±6.6%) and damaged cells (5.5±2.3 vs. 8.0±3.1%) was higher under aerobic conditions, but the percentage of uncertain material was higher under anaerobic conditions (20.4±5.8 vs. 17.1±3.2%) (Figure 7). None of these differences reached statistical significance. Overall, the aerobic processing of the stool sample leads to a slight decrease in the percentage of living cells compared to the semi-anaerobic procedure. This fact should be taken into account when evaluating the results.

**Figure 7:**
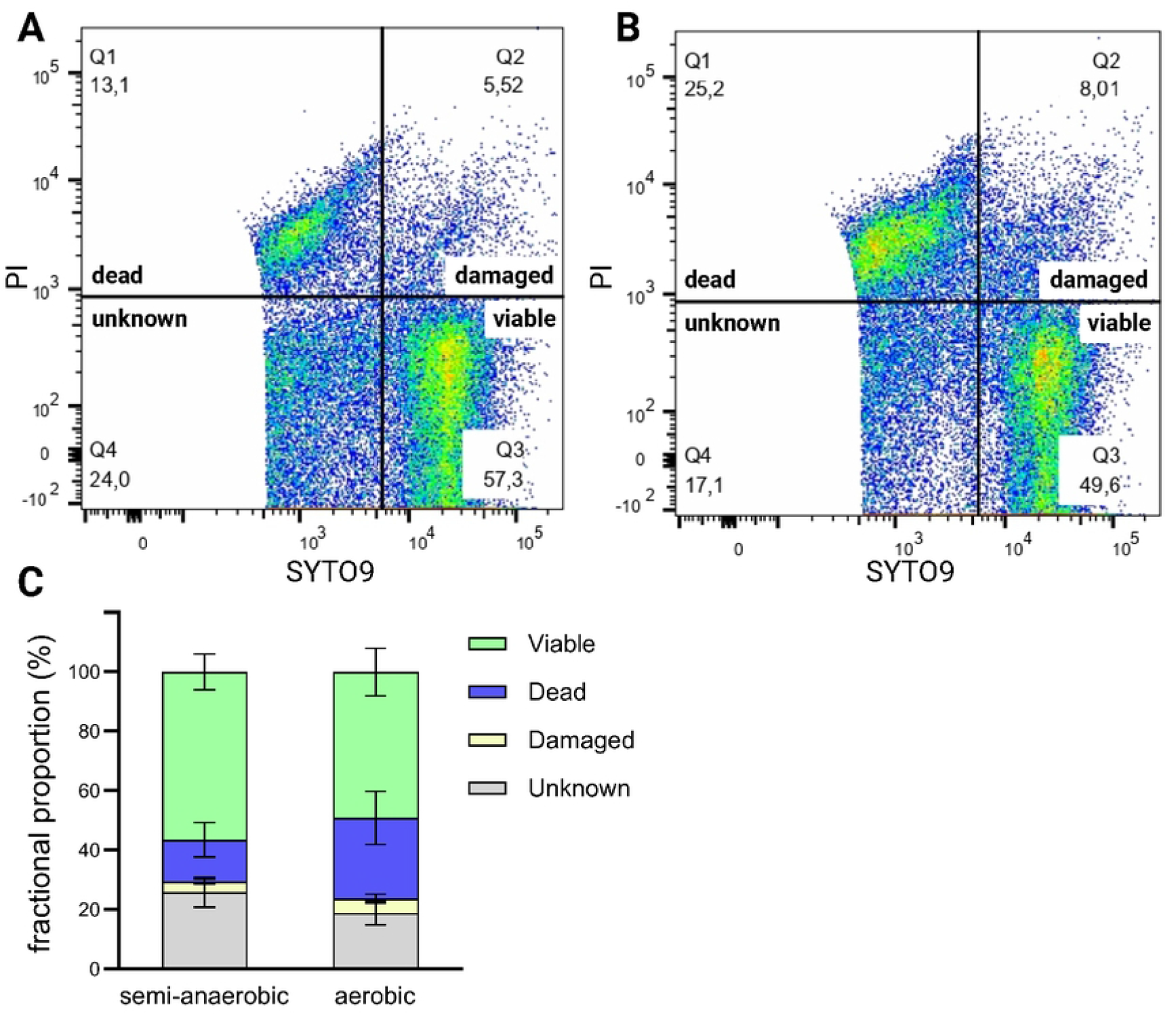
Quantification of the impact of aerobic and semi-anaerobic laboratory staining procedures on the viability of bacteria in human FMT stool filtrate sample. A: semi-anaerobic protocol; B: aerobic protocol; C: signal quantification.

### Viability in Donor FMT Filtrates

FMT filtrates stored for a year were analyzed for bacterial viability using both in vitro cultivation and membrane integrity methods. As for cultivation methods, notable differences in CFU counts were found between aerobic and anaerobic conditions. Importantly, we observed significant intrasample variability, with a median CV of 14.4% (min 4.1; max 41.2%) for aerobic cultivation and 17.0% (min 5.3; max 52.9%) for anaerobic cultivation. Flow cytometry outperformed cultivation in consistency, with a markedly lower median CV (0.9% for viable cells; max 8.5%, min 0.04%), underscoring its suitability for routine assessments (Tables 1 and 2).

**Table 1.** Number of CFU cultivated from frozen FMT filtrate stool samples in aerobic and anaerobic conditions. Values represents the means of technical triplicate ± s.d. Number of CFUs are normalized per 1 g of stool sample prior dilution.

| Donor ID | aerobic cultivation |  | anaerobic cultivation |  |
| --- | --- | --- | --- | --- |
| | CFU/g $\times 10^6$ | CV (%) | CFU/g $\times 10^8$ | CV (%) |
| 1 | $17.3 \pm 6.1$ | 35.1 | $7.1 \pm 2.9$ | 41.2 |
| 2 | $6.5 \pm 1.0$ | 15.0 | $22.2 \pm 7.4$ | 33.3 |
| 3 | $2.1 \pm 0.4$ | 18.3 | $0.2 \pm 0.03$ | 14.4 |
| 4 | $3.2 \pm 0.5$ | 16.4 | $2.1 \pm 0.1$ | 4.1 |
| 5 | $31.9 \pm 2.5$ | 7.8 | $1.3 \pm 0.2$ | 11.6 |
| 6 | $7.0 \pm 2.0$ | 28.9 | $0.7 \pm 0.1$ | 10.3 |
| 7 | $12.0 \pm 2.0$ | 17.0 | $4.6 \pm 1.1$ | 22.7 |
| 8 | $6.5 \pm 0.3$ | 5.3 | $4.2 \pm 1.3$ | 31.7 |
| 9 | $3.2 \pm 1.7$ | 52.9 | $2.3 \pm 0.3$ | 13.0 |
| 10 | $0.5 \pm 0.1$ | 19.1 | $5.9 \pm 1.4$ | 22.8 |
| 11 | 0.3 ± 0.01 | 14.1 | 6.7 ± 0.6 | 8.7 |
| 12 | 1.6 ± 0.4 | 24.7 | 10 ± 1 | 10.1 |
| 13 | 5.7 ± 0.9 | 15.1 | 6.3 ± 2.2 | 35.2 |

**Table 2.** The viability of bacteria in frozen FMT filtrate stool samples determined by flow cytometry. Values represents the means of technical triplicate ± s.d. and are given as the fraction representation in percent.

| sample ID | mean ± SD |  |  |  | CV (%) |  |  |  |
| --- | --- | --- | --- | --- | --- | --- | --- | --- |
|  | Viable | Dead | Damaged | Uncertain | Viable | Dead | Damaged | Uncertain |
| 1 | 36 ± 0.6 | 21.3 ± 1.3 | 5.7 ± 0.1 | 37.1 ± 0.7 | 1.7 | 6.1 | 1.3 | 1.8 |
| 2 | 36.3 ± 3.1 | 30.7 ± 2.9 | 7 ± 0.3 | 26.1 ± 0.1 | 8.5 | 9.4 | 4.1 | 0.2 |
| 3 | 24.2 ± 0.2 | 19.3 ± 0.1 | 7.1 ± 0 | 49.5 ± 0.1 | 0.8 | 0.3 | 0.4 | 0.2 |
| 4 | 29.8 ± 0.6 | 31.2 ± 0.5 | 6.5 ± 0.2 | 32.6 ± 0 | 2.2 | 1.8 | 2.3 | 0 |
| 5 | 20.8 ± 0.4 | 30.5 ± 0.1 | 4.5 ± 0.1 | 44.2 ± 0.2 | 1.9 | 0.3 | 1.6 | 0.5 |
| 6 | 26.3 ± 0.1 | 29.5 ± 0 | 5.2 ± 0 | 39.1 ± 0.1 | 0.6 | 0 | 0.9 | 0.1 |
| 7 | 65.9 ± 0 | 10.9 ± 0 | 4.2 ± 0.1 | 19.1 ± 0 | 0 | 0.5 | 1.6 | 0 |
| 8 | 41 ± 0.8 | 23.7 ± 0.4 | 9.2 ± 0.1 | 26.2 ± 0.6 | 2.1 | 1.5 | 0.8 | 2.3 |
| 9 | 43.6 ± 0.3 | 26.3 ± 0.8 | 6.7 ± 0 | 23.3 ± 0.5 | 0.7 | 3 | 0.4 | 2.1 |
| 10 | 43 ± 0.7 | 20.9 ± 0.5 | 3.7 ± 0.1 | 32.5 ± 0.1 | 1.5 | 2.6 | 3.9 | 0.2 |
| 11 | 45.6 ± 0.3 | 15.4 ± 0.1 | 6.6 ± 0 | 32.5 ± 0.2 | 0.5 | 0.6 | 0.2 | 0.5 |
| 12 | 44.5 ± 0.4 | 23.5 ± 0.5 | 7.4 ± 0.1 | 24.7 ± 0.2 | 0.9 | 2.1 | 0.7 | 0.6 |
| 13 | 38.8 ± 0 | 34.5 ± 0.6 | 9.5 ± 0.2 | 17.4 ± 0.4 | 0 | 1.6 | 2.4 | 2 |

## Discussion

In our investigation, we sought a method to accurately assess bacterial viability within faecal microbiota transplantation (FMT) suspensions. We optimized a commercially available kit, originally designed for pure bacterial cultures, to evaluate membrane integrity in stool samples. The main finding of this study is that this approach yields more reliable data than standard cultivation methods, making it preferable technique for evaluating the quality of the material for FMT.

Despite numerous stool banks and extensive research into FMT, standardized protocols remain elusive, with practices varying internationally. Some countries are moving to classify FMT filtrate as a pharmaceutical, prompting a push towards standardization (17). Stool, a biological substance, presents a complex challenge for characterization due to its varied components, including bacterial fragments and host metabolites (22,23). Crucial to FMT’s effectiveness is the count and variety of live bacteria present (8,15,16,24–28). However, consensus on a dependable method to quantify viable bacteria is lacking (20,21,29–32). FMT filtrate is inherently complex, containing diverse bacterial populations and other substances. Traditional cultivation methods are often considered the benchmark for assessing cell viability, yet they do not guarantee that non-culturable cells are non-viable (30,33,34). Culture-independent techniques, while bypassing culturability issues, have their own limitations (30). Fluorescent staining for membrane integrity, typically performed with flow cytometry, is a commonly adopted culture-independent method (35–37). Nevertheless, its effectiveness can vary across different bacterial taxa, and its application to the heterogeneous matrix of stool presents additional challenges (20,21,29,32)

Given these complexities, we compared the two prevalent methods—cultivation and membrane integrity assessment—for their accuracy and clinical applicability. Our adaptation of the LIVE/DEAD™ BacLight™ Bacterial Viability Kit for stool analysis demonstrated the potential to discriminate between microbial DNA signals and background noise effectively. We also noted that the type of stool processing, whether aerobic or anaerobic, could influence the viability results (16,18,29,38,39), with our study suggesting that aerobic conditions are adequate for routine viability assessments due to ease of handling.

Reproducibility is a crucial aspect of any laboratory method. Our study presents a significant step in addressing the variability associated with cultivation-dependent viability assessments. We observed high variability within samples using this method, with a mean CV of 17%. In contrast, the membrane integrity approach demonstrated remarkable consistency, with a median CV of 0.9% for viable cells.

Our research contributes to the field as the first to juxtapose these two viability assessment methods for feces. The lack of correlation between cultivation-dependent and membrane integrity results highlights the latter’s suitability for routine stool transplant quality assessments. The membrane integrity method combined with flow cytometry offers a consistent, reproducible, and efficient means to assess bacterial viability in stool grafts, without the need for specialized equipment or extensive technical expertise. We advocate for its inclusion in the standard evaluation of faecal material for FMT.

In conclusion, our findings recommend flow cytometry for evaluating bacterial viability in FMT, due to its accuracy and reliability over cultivation methods. This approach is likely to enhance the standardization of the quality control of faecal microbiota transplant preparations.

## Acknowledgement

We would like to thank all the volunteers who decided to participate in this study and research assistants, namely Šárka Gregorová, Nikola Bandíková, Šárka Vosalová, Kateřina Ťopková and Marie Chaloupecká for their valuable contributions to this project. We acknowledge also the Cytometry and Microscopy Facility at the Institute of Microbiology of the ASCR, v.v.i, Vídeňská 1083, Prague, CZ for the use of cytometry equipment and the support, namely to Jan Svoboda and Žaneta Slavíčková.

## Notes

### Competing Interest Statement

The authors have declared no competing interest.

